# Discovery of inhibitors of the *Pseudomonas aeruginosa* NADH:ubiquinone oxidoreductase (NQR) that hinder virulence factors

**DOI:** 10.64898/2026.02.25.708042

**Authors:** Sandaru M. Ileperuma, Jialin Li, Julian Ortiz, Ty Shade, Camille Vasseur, Natascia A. Ciancibello, Wael Elhenawy, Justin M. Di Trani

## Abstract

The growing threat of antimicrobial resistance has created an urgent need to identify novel therapeutic targets in bacteria. The NADH:ubiquinone oxidoreductase (NQR) is a potential target in a number of bacteria that transfers electrons from NADH to ubiquinone while pumping ions from the cytoplasm to the periplasm. In most species, this complex pumps sodium ions, whereas in *Pseudomonas aeruginosa* it pumps protons, thereby functioning as a member of the electron transport chain. Using a strain of PAO1 with the NQR knocked out, we demonstrate that the NQR complex plays a crucial role in the motility and biofilm development virulence factors in *P. aeruginosa*. We develop and execute a high-throughput inhibitor screen to identify and confirm compounds that inhibit NADH oxidation by this complex. Using single-particle cryogenic electron microscopy (cryoEM), we determine high-resolution structures of the NQR complex, both inhibitor-free and bound to one of the confirmed hits from our screen, demonstrating that it binds to the ubiquinone binding site. These structures provide insight into conformational dynamics controlled by binding at the ubiquinone site, with potential implications for the coupling between electron transfer and proton pumping in this complex. Biofilm development and motility assays with selected compounds from the screen show that they affect these virulence factors similarly to the NQR knockout.

## INTRODUCTION

Infections caused by antibiotic-resistant bacteria are projected to result in approximately 2 million deaths annually by 2050 (Naghavi et al., 2024), emphasizing the need for new targets and antibiotics. *Pseudomonas aeruginosa* is a Gram-negative bacterium with an exceptional ability to resist antibiotics (Pang et al., 2019) that infects immunocompromised individuals and causes chronic lung infections in those with cystic fibrosis (Lyczak et al., 2002). *P. aeruginosa’s* ability to establish infections and persist in the host is highly dependent on virulence factors such as biofilms. Biofilms are communities of bacteria encased in a self-produced matrix composed of DNA, polysaccharides, amyloids and other protein structures (Akbey and Andreasen, 2022). Biofilms allow *P. aeruginosa* to evade host immunity, outcompete other bacteria, and resist antibiotics (Flemming et al., 2016). While biofilms provide protection for *P. aeruginosa*, this environment is extremely demanding because nutrients and electron acceptors, such as oxygen, can be scarce. This demand is matched by the highly adaptable electron transport chain (ETC) of *P. aeruginosa*, which has drawn interest as a potential drug target (Borriello et al., 2004; Jo et al., 2022; Nguyen et al., 2011; Yoon et al., 2002). Motility is another key virulence factor in *P. aeruginosa* that relies heavily on the ETC. Flagellar rotation is driven directly by the proton motive force generated by the ETC, making it an energetically costly process (Doyle et al., 2004). Importantly, the ETCs of other bacteria (de Jager et al., 2020) and parasites (Baggish and Hill, 2002) have been successfully targeted. While the importance of the *P. aeruginosa* ETC for biofilm development, motility, and other functions that promote virulence is becoming apparent (Hazan et al., 2016; Jo et al., 2017; Raba et al., 2018; Xia et al., 2024), it is unclear which ETC components are crucial for these processes.

Components of the ETC transfer electrons from highly reductive substrates, such as NADH or succinate, to terminal electron acceptors, such as oxygen, while translocating protons from the cytoplasm to the periplasm of bacteria. In the canonical ETC, NADH and succinate are oxidized by complex I or complex II while reducing membrane-bound ubiquinone to ubiquinol. Electrons from ubiquinol are used by complex III to reduce cytochrome *c*, which ultimately reduces complex IV, allowing it to reduce oxygen into water. The *P. aeruginosa* genome codes for three enzymes that transfer electrons from NADH to ubiquinone. Complex I, the type II NADH dehydrogenase (NDH-2), as well as the NADH:ubiquinone oxidoreductase (NQR) complex. While in most bacteria, including *Vibrio cholerae*, the NQR complex pumps sodium ions, in *P. aeruginosa* this complex is a proton pump (Raba et al., 2018), making it a member of the ETC. It has been suggested that this complex is important for *P. aeruginosa’s* virulence (Hu et al., 2024; Raba et al., 2018) and is expressed under conditions mimicking infection (Liang et al., 2020). Importantly, NQR complexes have recently attracted considerable interest as drug targets in several bacteria, including *Vibrio cholerae, Vibrio alginolyticus* and *Chlamydia trachomatis* (Dibrov et al., 2017; González-Montalvo et al., 2025; Hu et al., 2024). However, there are very few known inhibitors of NQR complexes, with only eight reported (González-Montalvo et al., 2025). Interestingly, the NQR complex of *P. aeruginosa* (*pa*NQR) is resistant to a prominent inhibitor of the *V. cholerae* NQR (*vc*NQR), 2-heptyl-4-hydroxyquinoline N-oxide (HQNO) (Hau et al., 2023; Raba et al., 2018). HQNO is a quinolone analogue produced in *P. aeruginosa* that promotes the formation of biofilms by inhibiting complex III and creating reactive oxygen species that trigger cell autolysis (Hazan et al., 2016).

The best-characterized NQR complex is from *V. cholerae*, with several published cryoEM and crystallographic structures (Ishikawa-Fukuda et al., 2025; Kishikawa et al., 2022; Steuber et al., 2014). The *vc*NQR complex has six subunits, NqrA-F, with all subunits containing at least one transmembrane helix except for NqrA. The NqrF subunit harbors the riboflavin derivative, flavine adenine dinucleotide (FAD), where NADH oxidation occurs. Riboflavin derivatives are typical entry points for electrons in NADH oxidases (Liang et al., 2025;Lokendra K. Sharma et al., 2009). From the FAD at NqrF, the electron travels through a series of redox carriers, including other riboflavin derivatives, and eventually to the ubiquinone binding site in the NqrB subunit. Recent work identified the sodium-pumping mechanism of the *vc*NQR complex as the channel shared between the transmembrane domains, NqrD and NqrE (Ishikawa-Fukuda et al., 2026). The structural basis of proton specificity in the *pa*NQR complex remains poorly understood, as no structures of this complex are currently available.

In this work, we develop and implement a high-throughput enzymatic screen allowing us to discover small-molecule inhibitors of the *pa*NQR complex. Endogenous purification of the *pa*NQR complex allows us to obtain high-resolution structures using cryogenic electron microscopy (cryoEM); inhibitor-free and bound to one of the discovered compound hits, L-798106. These structures reveal inhibitor-induced changes in the complex’s structural dynamics and enable elucidation of ion-channel differences between *pa*NQR and *vc*NQR that may afford their ion specificity. Additionally, the L-798106-bound data reveals the structural basis for its binding. Excitingly, L-798106, as well as another inhibitor identified in the screen, appear to hinder *P. aeruginosa* biofilm development and inhibit swarming motility in this bacterium.

## RESULTS

### Discovery of compounds that inhibit the *pa*NQR complex

There are a small number of known inhibitors of the *vc*NQR complex (Dibrov et al., 2017, p.; González-Montalvo et al., 2025), and to our knowledge, none for the *pa*NQR complex. To identify inhibitors of the *pa*NQR complex, we developed a two-step series of NADH consumption assays. The first step was performed with crude membranes from *P*. aeruginosa with the *pa*NQR complex, and the second step was performed with *P*. aeruginosa membranes that were enriched for this complex (Fig. 1*A*), allowing us to confirm the hits. We tested both the LOPAC® library of pharmacologically active compounds and the ENAMINE Bioreference library, containing 1,280 and 2,405 compounds, respectively. The entire contents of both libraries were tested in the first screening step, and hits were confirmed with the second assay.

**Figure 1.**
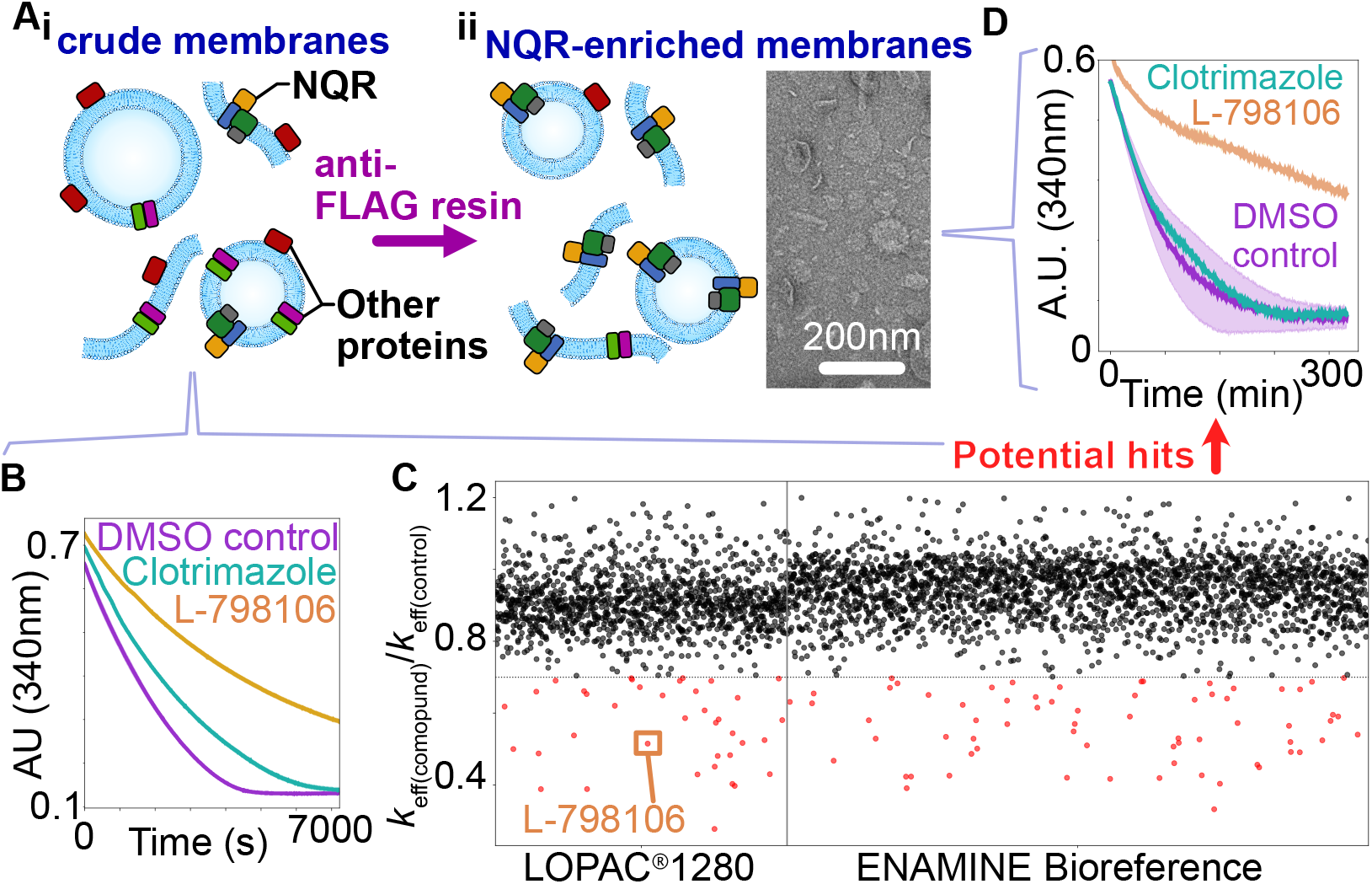
High throughput screening of compound libraries. (A) Schematic showing how crude membranes (i) extracted from *P. aeruginosa* and enriched for the NQR complex (ii - left), which were imaged using negative stain EM (ii -right). (B) Example NADH consumption assays for the first step of the screen with potential hits: L-798106 (orange), Clotrimazole (teal), and DMSO control (purple). (C) *k*_eff(compound)_/*k*_eff(control)_ for each compound in the LOPAC 1280 (left) and ENAMINE Bioreference (right) library, with the lowest 2% compounds shown in red below the horizontal threshold line. (D) Example NADH consumption assays for the second step of the screen with the same potential hits from B.

In the first screening step, we tested compounds for inhibition of NADH oxidation in a 384-well format using crude membranes prepared from *P. aeruginosa* (Fig. 1*Ai*). The presence of the *pa*NQR complex in the membranes was confirmed by immunoblotting for the 3×FLAG tag, which was engineered on the C-terminal end of the NqrF subunit (PA2994), via homologous recombination (*SI Appendix*, Table S*1*). These membranes show excellent NADH consumption activity (Fig. 1*B* – *purple curve*). Many compounds appeared to slow down the NADH consumption rate of the crude membranes to different extents (Fig. 1*B* – *orange and teal curves*). Calculating the relative NADH consumption rate in the presence of each compound compared to a DMSO control (*k*_eff(compound)_/*k*_eff(control)_) made it possible to quantify the extent of inhibition by both the LOPAC® 1280 and ENAMINE Bioreference libraries compounds (Fig. 1*C*). Compounds with a relative effective rate below 0.7 (bottom 2%, 106 compounds) were selected as candidates for the second screening step (Fig. 1*C* – *red data points, SI Appendix*, Fig. S2). While inhibition is obvious, there is still residual activity (Fig. 1*B*), likely due to the presence of multiple NADH dehydrogenases in these crude membrane samples. For this same reason, it is impossible to determine whether these hits from this first screening step are inhibiting NQR, other NADH dehydrogenases, or downstream ETC components, causing a buildup of reduced ubiquinol and inhibition of the dehydrogenases and are therefore only “potential” hits.

To distinguish between inhibitors targeting *pa*NQR versus other members of the ETC, NADH consumption assays were performed with membranes enriched for *pa*NQR. Enrichment was performed as described previously (Di Trani et al., 2025), via affinity purification of membranes produced from the *pa*NQR:3×FLAG strain using anti-FLAG resin (Fig. 1*A*). By avoiding solubilization, we were able to test potential inhibitors of the *pa*NQR complex in its native lipid environment, which includes membrane bound ubiquinone, avoiding artifacts brought about by solubilization and using short-tailed ubiquinone analogues (King et al., 2009). Many of the potential hit compounds from the crude membrane assay displayed greater levels of inhibition with enriched membranes (Fig. 1*D* – *orange curve*), while others inhibited very little (Fig. 1*D* – *teal curve; SI Appendix*, Fig. S3). It is expected that inhibitors of the *pa*NQR would show greater levels of inhibition with membranes enriched for this complex, while inhibitors of other enzymes would display less. A wide variety of inhibitors showed activity against enriched membranes, including several marketed drugs and well-known ETC inhibitors (*SI Appendix*, Fig. S3). Structures of the compounds that displayed a relative rate less than 0.45 are shown in the supplemental information (*SI Appendix*, Fig. S4). Below this threshold there are 8 and 14 compounds from the LOPAC® 1280 and ENAMINE library, respectively (*SI Appendix*, Fig. S3-4). The *pa*NQR inhibitors were generally more hydrophobic (Median ClogP:5.5) than the remainder of the libraries (Median ClogP:2.4), which is unsurprising for membrane protein inhibitors. Several compounds with very similar structures (GW5074 and Benzdioarone) met this cutoff, confirming the reproducibility of the screen. Additionally, many of these hits share common features, including being uncharged and containing numerous polycyclic aromatic rings linked with alkyl chains. These similarities suggest a common inhibitory site, which we hypothesize is the quinone-binding site.

From these 22 *pa*NQR inhibitors, we selected the LOPAC® 1280 compound L-798106 for structural analysis and biological assays. This inhibitor was chosen because it was part of the group that shared the consistent features mentioned above (*SI Appendix*, Fig. S4), and its commercial availability. This compound is a well-characterized, potent EP3 receptor agonist (Gallant et al., 2002), but, to our knowledge, has not been shown to inhibit ETC activity. The IC_50_ for the *pa*NQR complex was measured at ∼8 µM with crude membrane samples (*SI Appendix*, Fig. S*5*). Assessment of the IC_50_ with crude membranes provides an estimate of inhibitor strength in a native environment. To gain mechanistic insight into the binding of L-798106 and to help confirm the fidelity of these high-throughput screens, we then used cryoEM to determine structural details of its binding to the *pa*NQR complex.

### High-resolution structures of inhibitor-free and -bound *pa*NQR

Crude membranes isolated from *P. aeruginosa* were solubilized with the detergent dodecyl maltoside (DDM), and the complex was isolated with anti-FLAG resin, then further purified by size exclusion chromatography. Purified *pa*NQR was either left inhibitor-free or incubated with the LOPAC® 1280 hit L-798106, before cryoEM grid preparation. Data collection and structural analysis yielded high-resolution *pa*NQR structures for both conditions (3.2Å and 3.0Å for inhibitor-free and with L-798106, respectively; Fig. 2*A*, *SI Appendix*, Fig. S*6*, Table S*2-3*). Both structures contain density for all six subunits; however, similar to previous structures of *vc*NQR (Ishikawa-Fukuda et al., 2025; Kishikawa et al., 2022b; Steuber et al., 2014a), density for the highly mobile, soluble region of NqrF is not apparent in the map. The architecture of *pa*NQR was similar to that of *vc*NQR, as suggested by their high sequence similarity (∼61%). There is density for DDM and a phosphatidylethanolamine (PE) in deep crevices at the interface between NqrB and both NqrD and NqrE in either structure (Fig. 2*A*, *SI Appendix*, Fig. S*6 – pink density and structures*).

**Figure 2.**
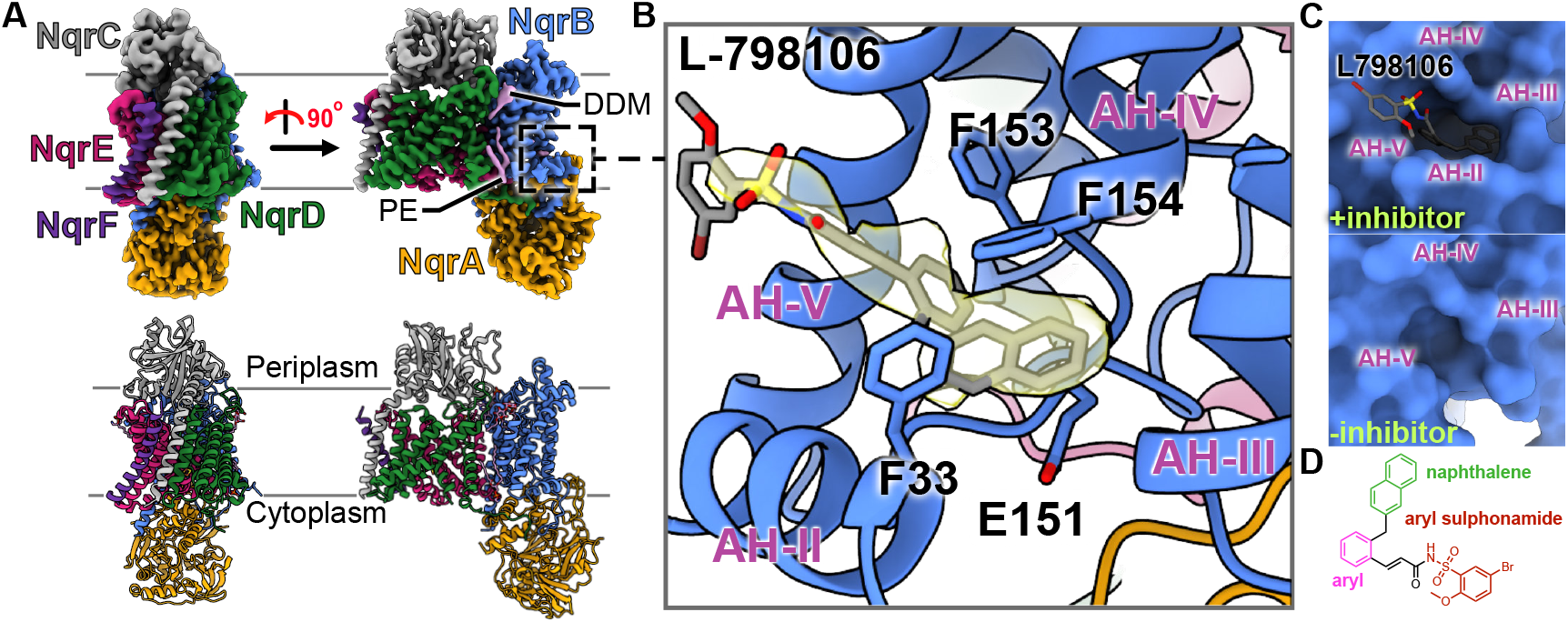
*pa*NQR bound to L-798106. (A) Overall cryoEM map (top) of the *pa*NQR and the atomic model (bottom). All subunits show clear densities except for the mobile cytoplasmic domain of NqrF. (B) Atomic model of the catalytic binding pocket of NqrB with L-798106. Residues of potential interest stabilizing the drug-bound form are labelled. (C) Surface model of the binding pocket with L-798106 (top), which stabilizes AH-II of NqrB, creating a deep cavity when compared to the apo structure (bottom). (D) Molecular structure of L-798106 with its main functional groups: naphthalene (green), aryl (pink) and aryl sulphonamide (red).

Density for L-798106 was found in the ubiquinone binding site located on the NqrB subunit, adjacent to the periplasmic side of the membrane and the NqrA subunit (Fig. 2*A*,*B*) (Kishikawa et al., 2022b). The binding pocket is composed of alpha helices (AH) III, IV and V, which are transmembrane helices and have strong density in both the inhibitor-free and L798106-bound structures, as well as AH-II on the N-terminal end of NqrB, which becomes ordered only in the bound structure (Fig. 2*B*, *SI Appendix*, Fig. S*7*). Together, these helices form a deep binding pocket that the naphthalene and aryl portion of L-798106 sit within (Fig. 2*B-D*). In the inhibitor-free structure, PHE154 appears to be in close contact with sections of the same peptide chain around GLU151 (*SI Appendix*, Fig. S*7*). Binding of inhibitor causes small movements in this region of NqrB, allowing F154 and the backbone portion E151 to pincer the naphthalene ring of L-798106, which would allow for F154 to engage in Pi stacking with the naphthalene ring (∼3Å, Fig. 2*B-D*; *SI Appendix*, Fig. S*7*) (Burley and Petsko, 1985). The naphthalene ring portion of L-798106 is also close enough to F33 on alpha helix II of NqrB to participate in a Pi-Pi interaction (∼3.5Å), this may be a major interaction locking this helix in place in the L-798106-bound structure. Another possible Pi stacking interaction occurs between F153 of alpha helix IV and the aryl group of L-798106, which are only ∼3.5Å apart. There are no obvious potential interactions between the NqrB and the aryl sulphonamide portion of L-798106. Furthermore, the density of the inhibitor in this region is less well defined, suggesting greater dynamics. The density for L-798106 in this map provides further confirmation of the success of this high-throughput screen and provides a structural basis for its binding.

In addition to the N-terminal end of NqrB, there appears to be ordering of the NqrA upon binding of L-798106. In the absence of an inhibitor, NqrA is highly mobile (Fig. 3*Ai*), bringing down its resolution in the nominal map (*SI Appendix*, Fig. S*6*). However, upon binding of L-798106, NqrA becomes ordered and tilts towards the binding site, reducing its mobility and increasing its local resolution (Fig. 3*Aii*, *SI Appendix*, Fig. S*6*). In addition to changes in mobility of NqrA, density for portions of NqrA and NqrB are much less well defined in the inhibitor-free structure, even after local refinement of this region (Fig. 3*B*, *SI Appendix*, Fig. S*6-7*). There is a complete lack of density for NqrA between the sections near G328 and F362 in the inhibitor-free structure, while with L-798106 this region appears ordered. Additionally, in the absence of an inhibitor, there is no obvious density for the N-terminal region of NqrB (anything N-terminal of N52), including the aforementioned alpha helix II, which is all ordered in the L-798106-bound structure. Together, this data suggests that binding of L-798106 causes structural stabilization of NqrA. Interestingly, in the L-798106-bound structure, the loop of NqrB between N52 and alpha helix II weaves through the NqrA subunit, making direct contacts with the region of NqrA between G328 and F362 (Fig. 3*B*, *SI Appendix*, Fig. S*7*). It’s possible that L798106 causes the N-terminus of NqrB to bind, rigidifying loops that weave through NqrA, limiting the dynamics of this subunit and interfering with its functionality.

**Figure 3.**
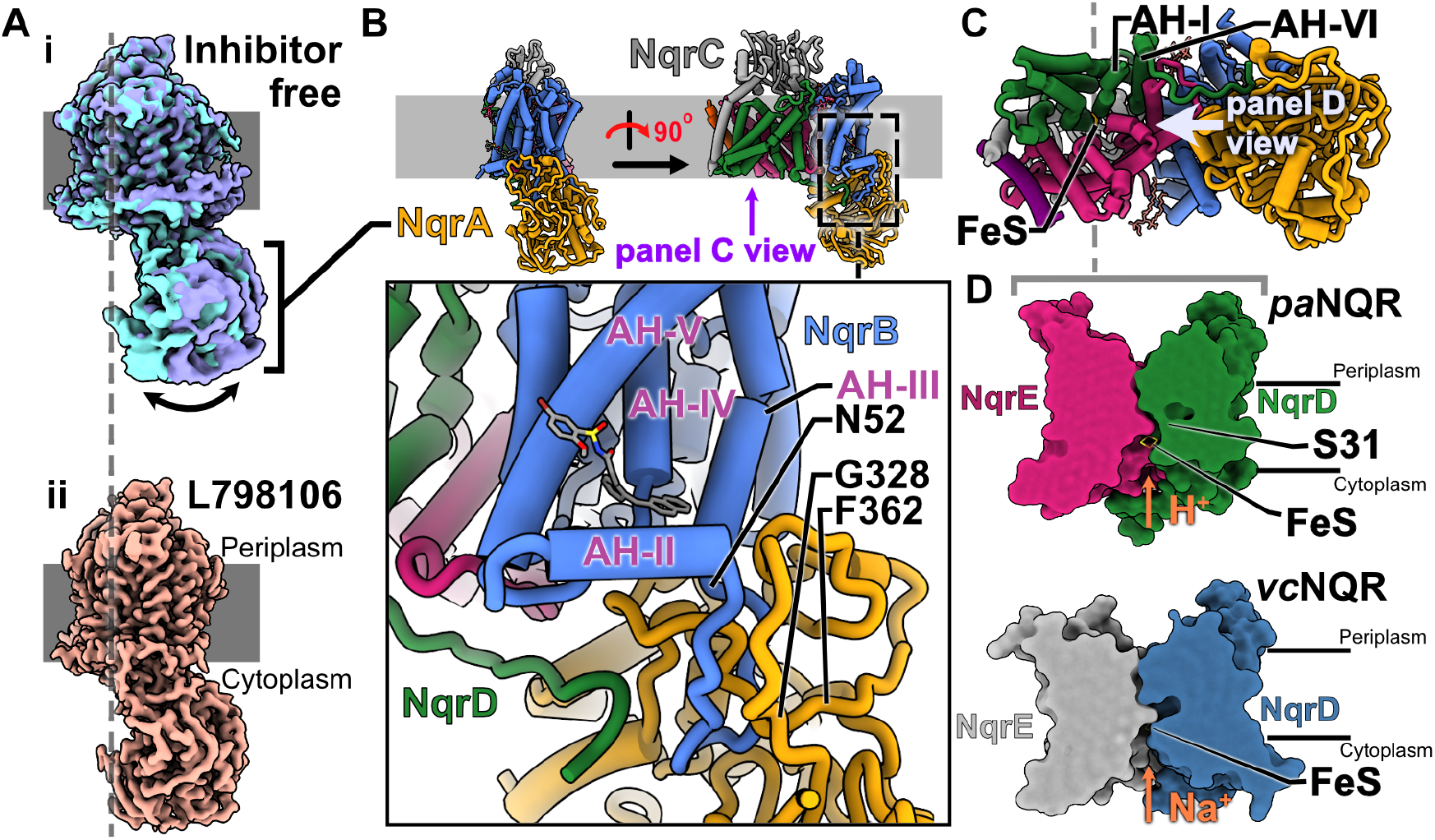
Structural dynamics of the *pa*NQR. (A)(i) Overlay of extreme points from 3D variability analysis for inhibitor-free dataset. (ii) nominal map of L-798106-bound dataset, which doesn’t reveal the same dynamics. (B) Ribbon structure showing interactions between NqrB and NqrA. (C) Top view of *pa*NQR. (D) Slice throughs of the proton channel formed between NqrE and NqrD in *pa*NQR (top) and *vc*NQR (bottom).

In addition to revealing changes in NqrA dynamics, these data enable a structural comparison of the *pa*NQR ion channel with previous *vc*NQR structures, providing insight into their ion specificity. In both the *pa*NQR and the *vc*NQR, NqrD and NqrE are arranged as a pseudodimer with a shared iron-sulphur cluster (Fig. 3*C*) forming the channel through which ions are pumped (Ishikawa-Fukuda et al., 2025). Several of the helices that make up this channel have different positions in the *pa*NQR and the *vc*NQR complex (RMSD=0.8, 0.85 for NqrE and NqrD, respectively; *SI Appendix*, Fig. S*8A*). However, the most considerable topological changes to the channel are caused by a shift in alpha helix I of the channel (Fig. 3*C-D*, *SI Appendix*, Fig. S*8A,B*). This shift causes significant changes in the shape of the channel in the area surrounding the iron sulphur cluster and alpha helix I, closing parts of the channel significantly in the *pa*NQR (Fig. 3*D*). This area of our inhibitor-free map is more diffuse than the L-798106-bound one, particularly around C31 of NqrD, which chelates the iron sulphur cluster (*SI Appendix*, Fig. S*8C*). The C-terminal loop extending from NqrD comes in contact with NqrA in the area of the inhibitor, forming a 6-7 amino acid interface with NqrA (Fig. 3*B*,*C*). Density for this NqrD loop only appears in the L-798106-bound structure, when NqrA is in the fixed position (*SI Appendix*, Fig. S7*B*). This loop extends from helix VI, which is directly adjacent to helix I of the proton channel (Fig. 3*C*). Interestingly, this loop is not present in *vc*NQR structures (Kishikawa et al., 2022c; Steuber et al., 2014b). The connection between NqrD and NqrA via this loop may allow for coupling between the NqrA subunit and the proton channel in the *pa*NQR.

### Testing *pa*NQR’s connection with virulence factors

Previous studies have shown that knocking out or inhibiting electron transport chain components can impair virulence factors, including biofilm development (Hazan et al., 2016; Jo et al., 2017; Sandoz et al., 2007; Xia et al., 2024). We set out to elucidate the dependence of planktonic growth, biofilm formation, and swarming motility on the *pa*NQR. This involved using a PAO1 mutant harboring a transposon insertion in the nqrB gene, which is expected to yield a catalytically inactive *pa*NQR complex (*nqrB::tn* strain). Additionally, we tested L-798106, and another confirmed hit from the ENAMINE Bioreference library, Zafirlukast, a leukotriene receptor antagonist.

First, planktonic growth was assessed by measuring optical density at 600nm in a 96-well plate with the wild type, knockout strain, and the addition of either inhibitor (Fig. 4*A*). These growth assays show no appreciable difference between conditions, suggesting that the *pa*NQR complex is not essential for planktonic growth. To explore the role of the *pa*NQR in biofilm formation, we used a crystal violet-based microtiter assay that stains extracellular polysaccharides, allowing quantification of biofilm production (Haney et al., 2021). The *nqrB::tn* strain showed a clear inability to form an effective biofilm in comparison to the wild-type PAO1 (Fig. 4*B*, *SI Appendix* Fig. S*9A*). L-798106 did not show an effect in decreasing biofilm, but Zafirlukast showed inhibition comparable to the *nqrB::tn* strain. We then decided to investigate compound effects on biofilm morphology using a Congo red-based staining assay (Jo et al., 2017), which stains amyloids in the biofilm, allowing for visualization of morphological features (Cabeen et al., 2016). The wild-type PAO1 strain showed a distinct wrinkling morphology, whereas the *nqrB::tn* strain showed a very smooth surface and was much less colored than the wild-type strain (Fig. 4*C*, *SI Appendix* Fig. S*9B*). This wrinkling of the colonies has been suggested as a marker of a more robust biofilm structure (Cabeen et al., 2016). Taken together with the crystal violet staining assay, it seems that removing *nqrB* diminishes the ability of PAO1 to form biofilms. Interestingly, the NqrB inhibitors, L-798106 and Zafirlukast, also interfered with biofilm formation, with fewer wrinkles formed by wild-type PAO1 in their presence compared to vehicle control. Interestingly, the colonies grown in the presence of L-798106 appear much darker than the other colonies, which may be due to the production of excess pigment molecules secreted by *P. aeruginosa* under different conditions (Abdelaziz et al., 2023). Finally, we assessed the effects of these inhibitors and the *nqrB::tn* strain on swarming motility assays, in which *P. aeruginosa* is applied to the center of a low-percentage agar plate containing minimal medium, allowing it to move outward in search of nutrients. In this assay, wild type outperformed all three of the other conditions, indicated by the size of the bacterial mass after 24 hours of incubation (Fig. 4*D*, *SI Appendix* Fig. S*9C*). Together, these data suggest that the *pa*NQR complex is important for the biofilm development and swarming motility virulence factors.

**Figure 4.**
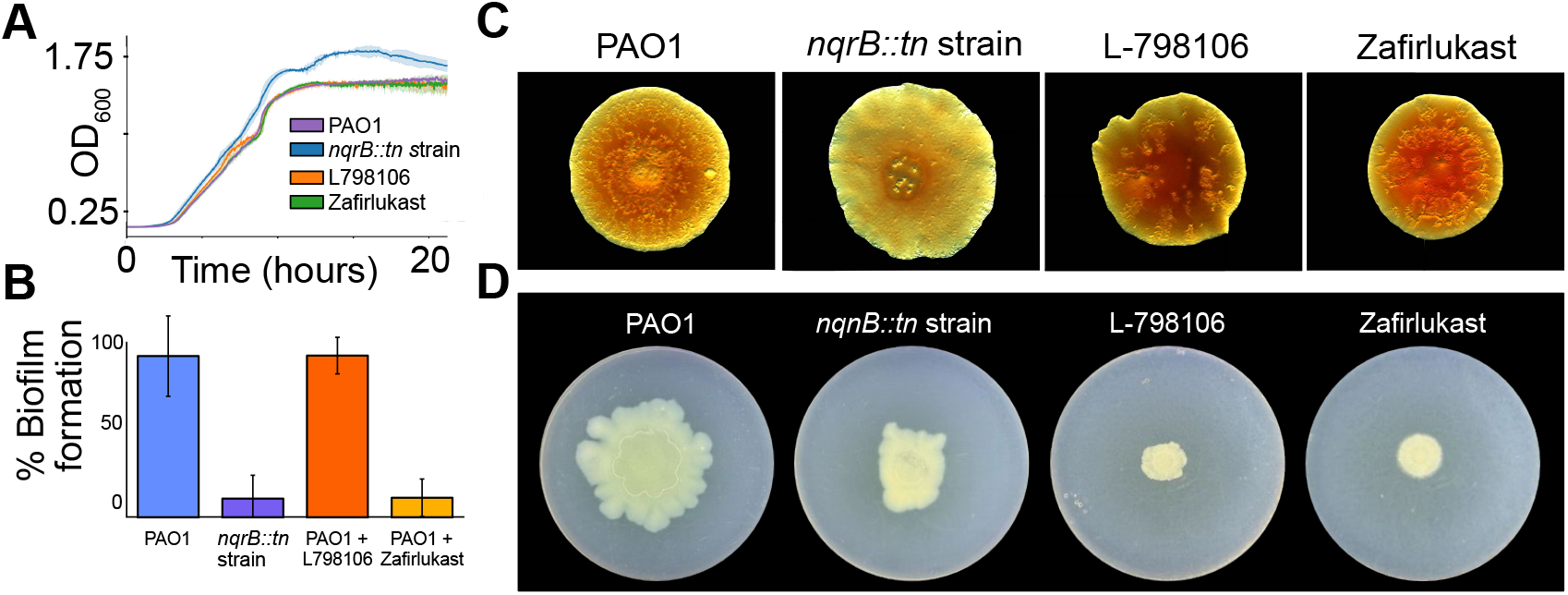
Biofilm development and swarming motility of *P. aeruginosa*. Wild type, ΔNqrB strains, L- 798106 and Zafirlukast were tested with (A) planktonic growth assays (B) biofilm formation assays (C) Colony morphology assays and (D) motility assays.

## DISCUSSION

This paper outlines the discovery of inhibitors of the *pa*NQR complex and provides structural insight into their binding mechanism. It also sheds light on the biophysical properties of this enzyme, including its dynamics and proton-pumping function. Importantly, these inhibitors and the knockout strain reveal the role of *pa*NQR in biofilm formation and motility.

Using a simple kinetic assay with crude membranes, we identify ∼100 potential inhibitors of the *pa*NQR complex and confirm many of them with an enriched membrane assay. The high NADH dehydrogenase activity in our crude membranes allows for over ∼10,000 kinetic traces to be measured with membranes isolated from just six litres of cells. The enriched membrane assays are also quite efficient, allowing for ∼1,000 kinetic traces with the same amount of starting material. This efficiency, combined with its speed, makes this series of kinetic screening assays a powerful method for discovering inhibitors of the *pa*NQR complex. In principle, these screening assays should be able to identify inhibitors of NQR complexes from other infectious bacteria and even different NADH dehydrogenases, highlighting the broader applications of this study. While L-798106 binds to the ubiquinone binding site, it is unclear where the other inhibitors bind without further analysis. The structural similarities among many of the inhibitors suggest they may bind to the same site; however, without further data, this remains unclear. Several of the inhibitors are dissimilar from L-798106 as they don’t contain a series of polyaromatic chains linked with alkyl chains. One of these outliers, diphenyleneiodonium chloride, showed potent inhibition in both crude and enriched membranes (*SI appendix*, Fig. S*2,3*). This compound is a known nonspecific inhibitor of flavoproteins (Lambert et al., 2008) and can inhibit *pa*NQR and other NADH oxidases by disrupting these sites, making it nonspecific. Another nonspecific compound we identified is sodium dodecyl sulphate, which disrupts membrane proteins. The discovery of these compounds from the screen and the presence of similar scaffolds among the hits demonstrate the assay’s fidelity.

The binding of L-798106 to the putative ubiquinone binding site induces ordering of sections of NqrA, NqrB and causes NqrA to become less dynamic. In these inhibitor-bound structures, the density for a loop connecting NqrD to NqrA is also visible. This structural insight is provocative as it may suggest a coupling mechanism between the binding site in NqrB and the proton channel. This model differs from the current understanding of how the *vc*NQR complex pumps protons, which is via electron transfer through the iron-sulphur cluster between NqrD and NqrE (Ishikawa-Fukuda et al., 2025). Unravelling these mechanisms will require more experimentation to prove. The C-terminal loop of NqrD is reminiscent of the lateral helix in complex I, which was suggested to couple redox reactions at the ubiquinone-binding site to the proton-pumping action in this enzyme (Kaila, 2018; Zhu et al., 2016). The structures presented here also enable comparison of the ion channels of the *vc*NQR and *pa*NQR complexes, providing a potential structural basis for ion selectivity in these enzymes.

The formation of biofilms and swarming motility are key virulence mechanisms deployed by *P. aeruginosa* to survive and evade antibiotics (Coleman et al., 2020; Hazan et al., 2016; Jo et al., 2017; Overhage et al., 2008). The *nqrB::tn* strain’s inability to form effective biofilms and its hindered swarming ability highlight the strong dependence of these virulence factors on this complex. Previous studies have demonstrated that HQNO is secreted by *P. aeruginosa* and promotes biofilm formation (Hazan et al., 2016). It’s been suggested that the lack of inhibitory activity by HQNO on the *pa*NQR may avoid autotoxicity (Raba et al., 2018) when secreting HQNO. This autotoxicity may explain why biofilm formation is hindered in the *nqrB::tn* strain in our assays (Fig. 4*B*,*C*). While Zafirlukast appears to inhibit biofilm development, L-798106 does not (Fig. 4*B*). Further, the colony morphology with these two compounds is less wrinkly than that of the wild type (Fig. 4*C*). This morphological feature has previously been shown to be linked with the redox state of *P. aeruginosa* (Dietrich et al., 2013). Furthermore, the *nqrB::tn* strain and both compounds show strong effects on motility, which is consistent with this being a less protective state for *P. aeruginosa*. While the exact mechanism by which these factors are hindered by the absence of the *pa*NQR remains unclear, these results highlight the clear importance of this enzyme for these virulence factors.

## MATERIALS AND METHODS

### Molecular and microbiological techniques, bacterial strains, plasmids, and growth conditions

Standard protocols were used for molecular and microbiological techniques (Sambrook, 2001). Genomic DNA isolation, plasmid preparation, and DNA gel extraction were performed using kits purchased from QIAGEN. All primers were obtained from Thermo Fisher Scientific.

All *P. aeruginosa* strains were derived from PAO1. *P. aeruginosa* mutant strains were generated using two-step allelic exchange, as described previously (Hmelo et al., 2015). A complete list of all bacterial strains and plasmids used in this study is found in *SI Appendix*, Table S1.

For the two-step allelic exchange, lysogeny broth (LB) contained 10.0 g tryptone, 5.0 g yeast extract, and 5.0 g NaCl per liter of ultrapure water. Vogel-Bonner minimal medium (VBMM) was prepared as a 10× concentrate, which contained 2.0 g MgSO_4_·7H_2_O, 20 g citric acid, 100 g K_2_HPO_4_, and 35 g NaNH_4_HPO_4_·4H_2_O, per litre of ultrapure water and was adjusted to pH 7.0 and sterilized by filtration. Semisolid media was prepared by adding 1.5% (w/v) agar to LB and VBMM. Where needed for selection, gentamicin was added to growth media at 60 μg/mL for *P. aeruginosa* and 10 μg/mL for *E. coli*.

### Construction of *P. aeruginosa* strain with a PA2994 C-terminal 3×FLAG sequence

Constructs of *PA2994* with sequence encoding a 3×FLAG sequence at the 3′ end of the *PA2994* gene were generated from *P. aeruginosa* PAO1 with a protocol described previously (Hmelo et al., 2015). Briefly, the pEX18Gm plasmid was linearized with forward (5’-GAGGATCCCCGGGTACCGA-3’) and reverse (5’-TAGAGTCGACCTGCAGGCAT-3’) primers. This linearized plasmid was ligated via Gibson assembly to a DNA strand containing an 800bp region of homology surrounding the 3×FLAG amino acid at the 3’ end of *PA2994*. The resulting allelic exchange vector, pEX18Gm::PAJDT2 was used to transform *E. coli* strain SM10(λpir) and selected for on LB agar supplemented with gentamycin, identified by colony PCR, and verified by Sanger sequencing using M13 forward and M13 reverse primers.

The insertion allele encoded by pEX18Gm::PAJDT2 was introduced into *P. aeruginosa* PAO1 via biparental mating with the donor strain *E. coli* SM10 (de Lorenzo and Timmis, 1994; Hoang et al., 1998; Thöming et al., 2020). Merodiploids were selected on VBMM supplemented with 60 μg/mL Gen. SacB-mediated counter-selection was carried out by selecting for double cross-over mutations on no-salt LB (NSLB) agar supplemented with 15% (w/v) sucrose. 3×FLAG insertion was identified by colony PCR with primers flanking the outside regions of the upstream and downstream regions of the sequence. PCR products were gel-purified and sent for Sanger sequencing to confirm insertion.

### Western blot analysis

PAO1 3×FLAG-tagged NQR bacterial cultures were grown in LB or TB, for 24 h at 37 °C with shaking at 200 rpm. Cells were harvested by centrifugation at 4,000 rpm for 10 min at 4 °C. Pellets were resuspended in lysis buffer (200 mM NaCl, 200 mM Tris–HCl pH 7.5 at 4ºC, 5 mM EDTA, 1% Triton X-100, 1 mM PMSF, 1 mg.mL⁻1 DNase, protease inhibitor cocktail) at a 1:3 pellet-to-buffer ratio and incubated for 1h at 4 °C on a nutator. Lysates were clarified by centrifugation at 15,000 rpm for 30 mins at 4 °C. Supernatants were mixed with 2× loading dye, and 30 µL of each sample were loaded on SDSPAGE 12% polyacrylamide gels and transferred onto nitrocellulose membranes (0.45 µm; Bio-Rad) at 30 V overnight at 4 °C. Membranes were blocked in TBST (Tris-buffered saline containing 0.1% Tween) supplemented by 5% BSA for 1 h, then incubated with anti-FLAG M2 mouse monoclonal antibody (1:15,000) for 1 h at room temperature, followed by Anti-Mouse IgG antibodies (1:15,000). Between each antibody, the membrane was washed three times with TBST Buffer. Proteins were detected using enhanced chemiluminescence (ECL; Cytiva) and imaged with a ImageQuant LAS 4000 imaging system (GE HealthCare). Unprocessed blot images are provided in the Supplementary *Information*.

### Growth and purification

*P. aeruginosa* was cultured in LB medium. A preculture in liquid medium (∼20 mL) was inoculated with cells from an agar plate and grown at 37 °C with shaking at 180 rpm overnight. This culture was used to inoculate a larger culture (6 L), which was grown under the same conditions in Fernbach flasks for an additional 24 h. Cells were harvested by centrifugation at 4,800 ×g for 20 min and resuspended in lysis buffer (5 mL per 1 L of culture; 0.2 M Tri-HCl pH 8, 1 M sucrose, 1 mM EDTA, 1 mg/mL lysozyme), incubated at room temperature (RT) for 5 min, and 20mL of water was added for every 5 mL of lysis buffer. The resulting mixture was swirled gently, placed on ice for 20 min, and centrifuged at ∼200,000 ×g for 45 minutes at 4 °C. The pellet was resuspended in homogenization buffer (50 mL per 1 L of culture; 10 mM Tris-HCl pH 7.5, 5 mM EDTA, 1 mM 1,4-dithiothreitol, 7 μg/mL DNase I). Cells were then broken by two passes through a continuous flow cell disruptor (Avestin) at 17 kpsi. Cell debris was spun down by centrifugation at 10000×g at 4 °C, and the supernatant was centrifuged at 200,000 ×g for 1 hour at 4 °C to harvest membranes. Membranes were gently resuspended in solubilization buffer (50 mM Tris-HCl, pH 7.5, 0.5 mM EDTA, 10% (v/v) glycerol) to a concentration of 3 mg/mL total protein, flash frozen in liquid nitrogen, and stored at -80 °C until use. For purification, the membranes were thawed and solubilized by gentle tumbling with 1% (w/v) n-Dodecyl-β-D-Maltopyranoside (DDM, Anatrace) for 45 min. Insoluble material was removed by centrifugation at 180,000 ×g for 50 min at 4 °C. Solubilized protein was loaded onto 0.5-1.5 mL of M2 resin (Sigma), which was equilibrated with DTBS buffer (50 mM Tris-HCl, pH 7.4, 150 mM NaCl, 0.02 % (w/v) DDM). The column was then washed with ∼5 column volumes of DTBS before elution with ∼8 column volumes of 150 μg/mL 3×FLAG peptide in DTBS.

### NADH consumption assays for screening compounds and measuring IC_50_’s

1μl LOPAC® 1280 compounds were pipetted into a 384 well plates from 1mM stocks. 0.09ul of the Enamine bioreference compounds were dispensed using an ECHO 650 acoustic dispensing robot from 10mM stocks into 384-well plates. Crude membranes from *P. aeruginosa* grown in LB media were isolated as above. Membranes were clarified with a 6,000 ×g spin down for 10 minutes at 4°C. The supernatant was carefully diluted 30-fold into TBS buffer. 30 µL of the diluted membrane was then injected into each well of the 384-well plate already containing compounds using the BioTek Synergy Neo2 plate reader injection system. The plates were incubated for 1 hour at 37 °C in the plate reader, after which 30 μL of 300 μM NADH was injected into the wells, bringing the total reaction volume to 60 μL. Decrease in absorbance readings at 340nm was observed over the course of the reaction and compared with the DMSO controls, 1.7% and 0.15% for the LOPAC Enamine Bioreference library, respectively.

For enriched membrane NADH consumption assays, crude membranes were spun down at 4,000 ×g for 10min at 4 °C, then filtered using a 0.45 μm sterile filter and then applied to an M2 resin column. The column was washed with TBS and then eluted with 150 μg/mL 3×FLAG peptide in TBS before concentrating using a 100 kDA filter. Per 2L of membranes, the elution was diluted to give a total volume of 7.5ml in TBS. 29ul was then injected into a 384-well plate containing 1ul of compounds from 1mM stocks, yielding a final concentration of 16.7uM. 30 ul of 300 μM NADH in TBS was then injected. Readings were taken at 340nm over 6 hours and compared with 1.7% DMSO controls.

Briefly, NADH consumption curves were fit to negative exponentials to extract effective rates. The effective rate in the presence of each compound was divided by the effective rate of the vehicle control in the same row of the well plate to calculate the relative rate. This was performed with both the crude membranes and the NQR-enriched membranes.

### IC_50_ of L-798106

Membranes prepared as outlined previously were spun down at 6,000g for 10 minutes at 4°C to remove cellular debris. 384 well plates were dispensed using the ECHO 650 acoustic dispensing robot to have 600nL of 10^−2^ mM, 10^−3^ mM, 10^−4^ mM, 10^−5^ mM, 10^−6^ mM, 10^−7^ mM of the desired compounds. 30 μL of the membranes were injected into the well and incubated at 37°C for 1 hour followed by injection of 30 μL of 300 μM NADH disolved in TBS. This gave a final concentration gradient of 100 μM, 10 μM, 1 μM, 0.1 μM, 0.01 μM and 0.001 μM. Absorbance readings at 340nm were taken throughout the course of the reaction. The readings were then compared to the 1% DMSO control. The IC_50_ of L-798106 was determined by measuring the effective rate at varying inhibitor concentrations and fitting the data to a sigmoidal function.

### Negative stain EM of enriched membranes

Crude membrane preparations were diluted 1 in 60. 4µL of this was dispensed into a glow-discharged 400 mesh Cu-Rh grid with a 20nm continuous carbon film. The excess sample was blotted using filter paper and then washed with MilliQ H_2_O and dried with another filter paper. 3.5µL of UAR-EMS Uranyl Acetate replacement stain was applied to the grid. Excess stain was blotted onto filter paper and incubated for 15 minutes at room temperature, then air-dried. Images were collected using a Talos L120C.

### Grid preparation and EM data collection

For cryoEM grid samples, gel filtration was done using DTBS buffer (50 mM Tris-HCl, pH 7.4, 150 mM NaCl, 0.02 % (w/v) DDM) on the resin-purified sample using a superose™ 6 Increase 10/300 GL. Gel filtration fractions containing *pa*NQR were pooled and concentrated to ∼8 mg/mL using a 0.5 mL concentrator with a 100 kDa MWCO (Vivaspin, Sartorius). The sample was either left in the absence of inhibitor or incubated with 100 μM of L-798106, and 2 μL was applied to nanofabricated holey gold grids (Marr et al., 2014) that were glow-discharged for 2 min in air (20 mA with a PELCO easiGlow) immediately before use. Grids were blotted for 3.5 s with an EM GP2 Plunge Freezing device (Leica Microsystems) before freezing in liquid ethane. CryoEM data were collected with a Titan Krios G3 (inhibitor-free) or G4 (with L-798106) electron microscope (Thermo Fisher Scientific) operated at an extraction voltage of 300 kV and equipped with a Falcon 4i direct detector camera, and Selectris X energy filter. The data collected was using the EPU software package. For the inhibitor-free sample, a dataset of 8,388 EER movies was collected, with each movie consisting of 40 exposure fractions over ∼4.3 s, a total exposure of ∼40 e^-^/Å^2^, exposure rate of ∼10.7 e^-^/pixel/s and a magnification of 130,000× with a calibrated pixel size of 0.93 Å. For the L-798106-bound dataset 12,408 TIFF movies were collected, with each movie consisting of 40 exposure fractions over ∼4 s, a total exposure of ∼39 e^-^/Å^2^, exposure rate of ∼9 e^-^/pixel/s and a magnification of 130,000× with a calibrated pixel size of 0.94 Å (*SI Appendix*, Table *S3*).

### CryoEM image processing

Image analysis was performed using cryoSPARC v4.7.1 (Punjani et al., 2017), with all refinements applied as non-uniform refinements (Punjani et al., 2020). Collected movies were aligned with patch motion correction, and CTF parameters were estimated with patch CTF. The CTF parameters, along with manual curation, were used to discard movies of low quality. This led to a final data set of 8,038 and 9,826 micrographs for the inhibitor-free and L-798106 dataset, respectively. Ab-initio followed by and heterogeneous refinements were used iteratively to clean out “junk particles”. The final particle set were then extracted with a box size of 288 pixels. This resulted in 79,904 particles for the inhibitor-free *pa*NQR dataset and 452,841 for the L-798106 dataset. 3D variability analysis (Punjani and Fleet, 20210) with a filter resolution of 8 Å and a mask around the NqrA subunit was used to reveal its rocking motion. Local refinement was performed on the inhibitor-free dataset with masks either around NqrA or NqrB-F.

### Model building and refinement

The initial model was built using the L-798106 map using a rigid body fitting of the alpha-fold model (Jumper et al., 2021) of the subunits in Coot v0.9.8.96 (Emsley et al., 2010). This was then refined using ISOLDE v1.10.1 (Croll, 2018) and Coot iteratively. L-798106 was parameterized in ISOLDE and the MD force field enables better fitting of the compound within the density after rigid body fitting in coot. This model was then used to build the inhibitor-free *pa*NQR structure using the same process. The final refinement was done using PHENIX v1.21.2-5419 (Liebschner et al., 2019). All figures were made and rendered using UCSF ChimeraX (Pettersen et al., 2021).

### Biofilm production assay (crystal violet staining)

1×M63 media was prepared containing 2g/L (NH_4_)_2_PO_4_, 13.6g/L K_2_HPO_4_, 0.5 mg/L FeSO_4,_ and supplemented to be 0.2% glucose, 0.5% casamino acid and 1mM MgSO_4_. Pre-cultures were grown overnight in LB and standardized to an OD of 0.025 in a 5ml inoculation test tube containing 2ml of 1×M63 media. Compounds were added to a final concentration of 15µM and 0.15% DMSO. Test tubes were placed in the incubator overnight at 37ºC at 200 rpm for 24 hours. The solution in the test tubes was then discarded, and biofilms were washed with MilliQ water. The adhered biomass is quantified by staining with 2.5 ml of 1.7% crystal violet and vortexed. After discarding the excess stain, the tubes were once again washed with MilliQ water and then the adhered biofilm was solubilized with 2mL of a 80% ethanol/20% acetone solution. OD_595nm_ was measured to quantify biofilm production.

### Biofilm development assays

2×M63 media was prepared containing 100mM KH_2_PO_4_, 15mM (NH_4_)_2_SO_4_, 0.5mg/L FeSO_4_ · 7H_2_O and supplemented with 1% casamino acid, 0.4% glucose and 2mM MgSO_4_. 3% agar was prepared by dissolving 3g of agar in 100mL of MilliQ H_2_O. Congo red dissolved in MilliQ was added to the 2×M63 media to a final concentration of 80μg/mL. Compounds were directly added to the media from 10-20mM stocks at 2× concentrations into the 2× M63 media to keep DMSO between 0.15-0.3%. The resulting media was kept warm in a hot water bath at 50°C. The warmed 3% agar was then mixed 1:1 with the 2× M63 media to give final concentrations of 15μM compounds, 0.3% DMSO, 40μg/mL congo red, 1.5% agar and 1×M63 media. 4mL of M63 agar media was poured into 6-well plates. PAO1 wt and ΔNqrB strains were grown for 16-18 h in LB media at 37°C. OD_600_ of both cultures was normalized to 1. 2μL of standardized inocula were spotted onto the center of the media carefully so as not to pierce the agar medium. Plates were incubated at 37°C for 4 days with images taken every 24 hours using a Leica M125C microscope attached with a Nikon DS-Fi1 camera.

### Swarming Motility assay

The motility assay was conducted according to procedures outlined by Ha et al (Ha et al., 2014). 5× M8 media containing: 30 g of Na_2_HPO_4_, 15 g KH_2_PO_4_ and 2.5g NaCl was diluted to 1× and supplemented with final concentrations of 0.5% casamino acid, 0.2% glucose, 1mM MgSO4 and 0.5% agar. Compounds were added to the warmed mixture to obtain a final concentration of 15µM and 0.15% DMSO. The mixture was gently swirled thoroughly to mix all the components, and thick plates were poured. Bacterial cultures grown overnight in LB media were standardized to an OD_600nm_ of 1. Plates were seeded with the standardized inoculum with 2µL to the center of the plate and incubated at 37°C for 24 hours prior to imaging.

## AUTHOR CONTRIBUTIONS

J.D.T. designed research; J.D.T performed genetic engineering; S.M.I. prepared NQR sample, collected and analyzed cryoEM data and performed and analyzed kinetics experiments with help from J.D.T; J.L. and T.S. helped with kinetics experiments and biological assays; S.M.I and J.O. performed biofilm experiments; J.D.T. supervised the cryogenic electron microscopy studies; W.E. designed and supervised biofilm experiments; J.D.T. and S.M.I. wrote the paper with suggestions from all other authors.

## Data Deposition

Data deposition: The electron cryomicroscopy map and associated models described in this article have been deposited in the Electron Microscopy Data Bank (EMDB) and the Protein Data Bank (PDB).

## ACKNOWLEDGEMENTS

This work was funded by the Cystic Fibrosis Canada 2024 Seed Grant, LKSIoV and University of Alberta Faculty of Medicine and Dentistry Startup funding (RES0067506 and RES0067539), Striving for Pandemic Preparedness – The Alberta Research Consortium (SPP-ARC, RES0069636). S.M.I was supported by a CAN-AMR Scholarship award and a LKSIoV Graduate Studies Entrance Award. N.A.C was supported by a CGS-M award. The Advanced Cell Exploration Core provided the LOPAC and ENAMINE libraries. The inhibitor-free cryoEM dataset was collected at the High-resolution high-throughput CryoEM facility at The Hospital for Sick Children, Toronto. The L-798106-bound cryoEM dataset was collected at the Alberta CryoEM Facility. We thank Dr. Samir Benlekbir and Dr. Zhijie Li at these facilities for their help with the collection. W.E. holds a Canada Research Chair in Microbiome Research (CRC-00249). W.E. is funded by Crohn’s & Colitis Canada (GAR 1034447), Natural Sciences and Engineering Research Council of Canada (DGECR-2023-00140), Canadian Institutes of Health Research (RES0069043), LKSIoV Research Support and Innovation Grant (RES0057474), and Striving for Pandemic Preparedness - The Alberta Research Consortium (RES0066221).

