## Supplemental Figures 1-9, Tables 1-3 for "Discovery of inhibitors of the *Pseudomonas aeruginosa* NADH:ubiquinone oxidoreductase (NQR) that hinder virulence factors"

**This PDF file includes:**

Supplementary Figures 1-9

Supplementary Tables 1-3

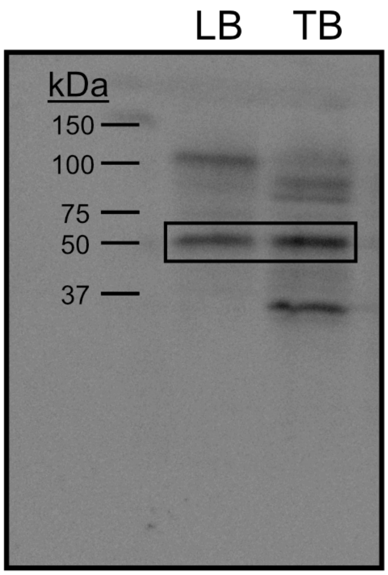


**Figure S1.** The ladder and its corresponding molecular weights are shown on the far left; the two conditions, LB and TB, are labelled accordingly, with the chemiluminescence band from NqrF highlighted within the box.

LOPAC Hits - Page 1

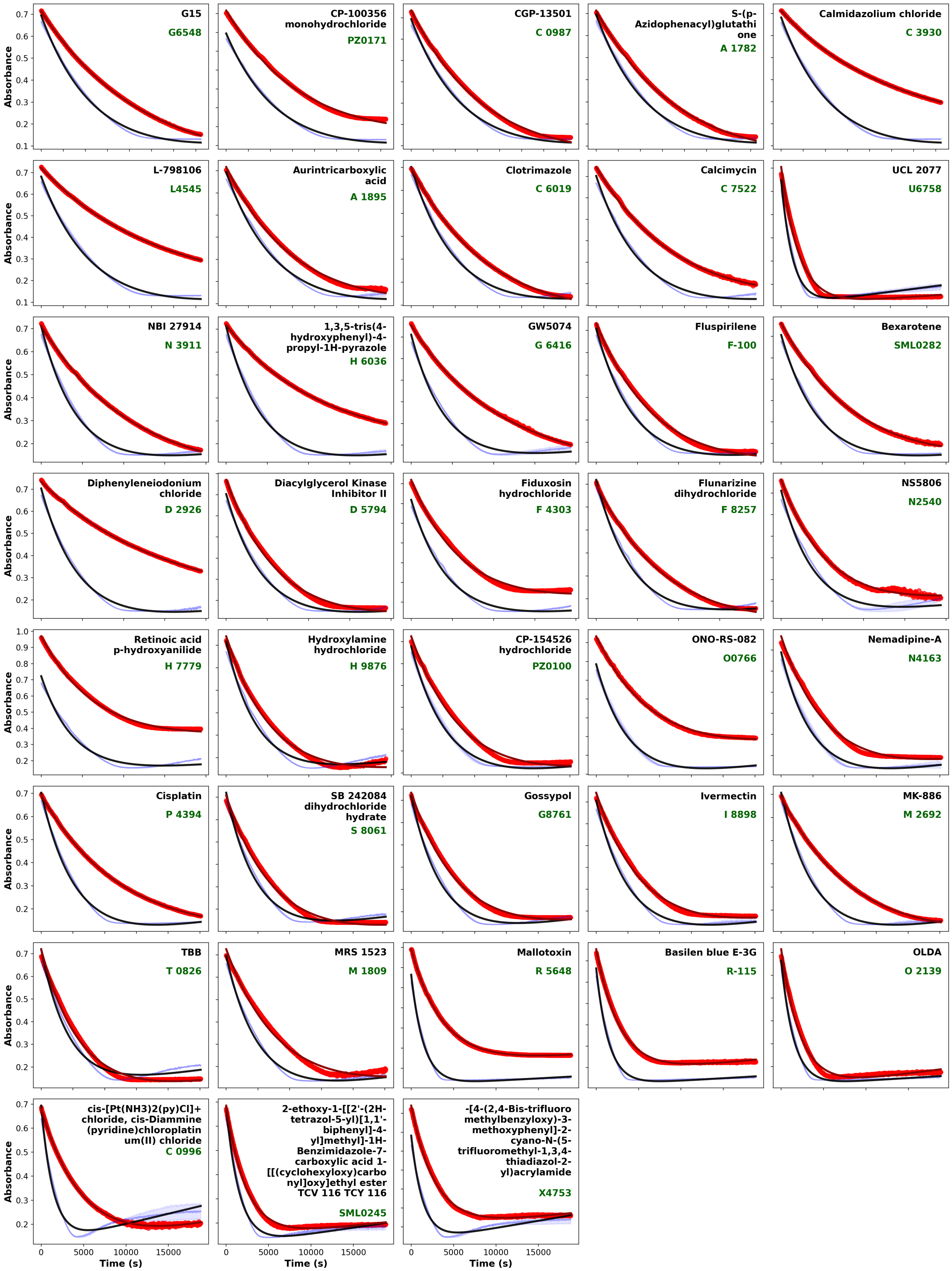

#### Enamine Hits - Page 1

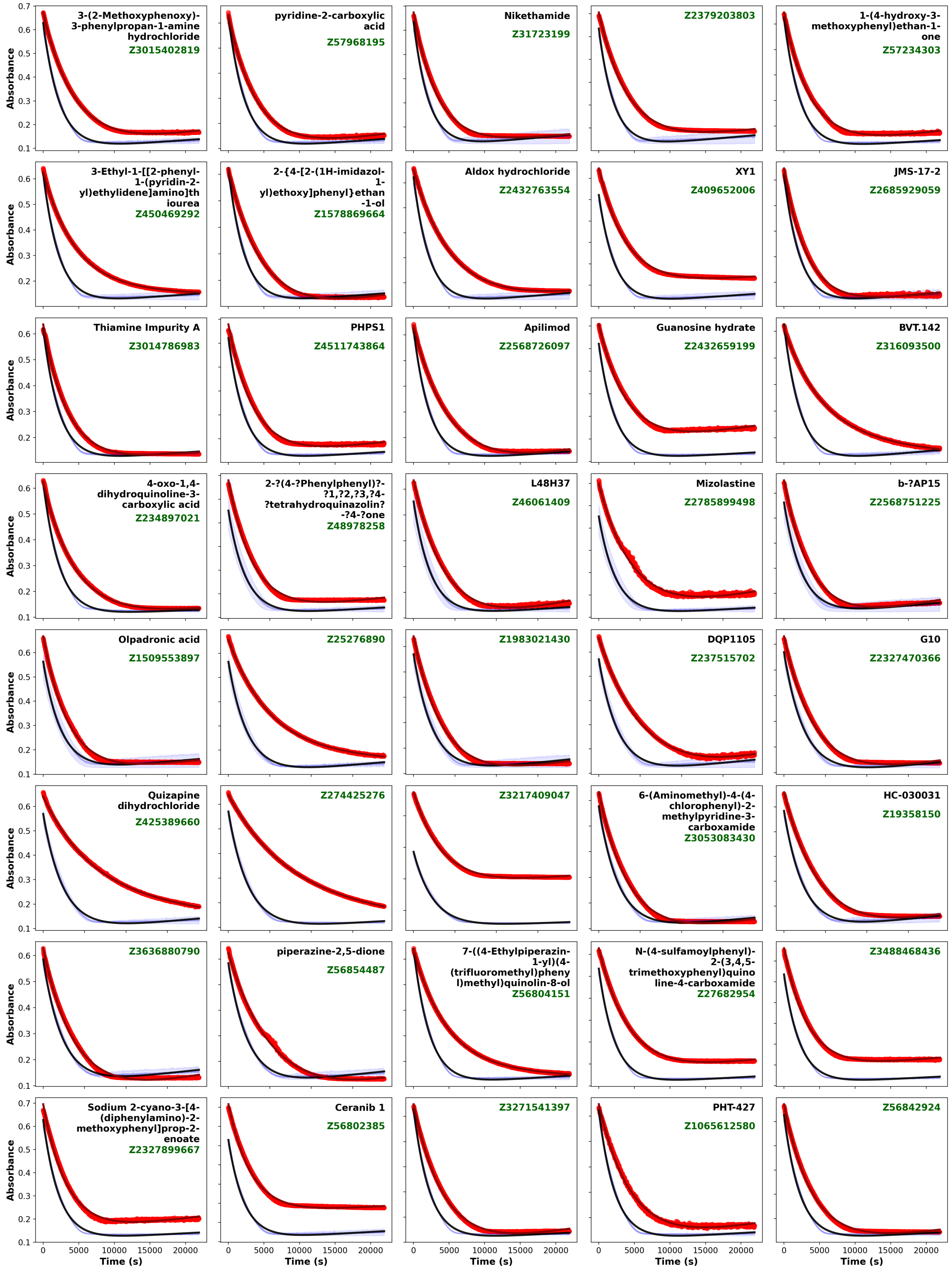

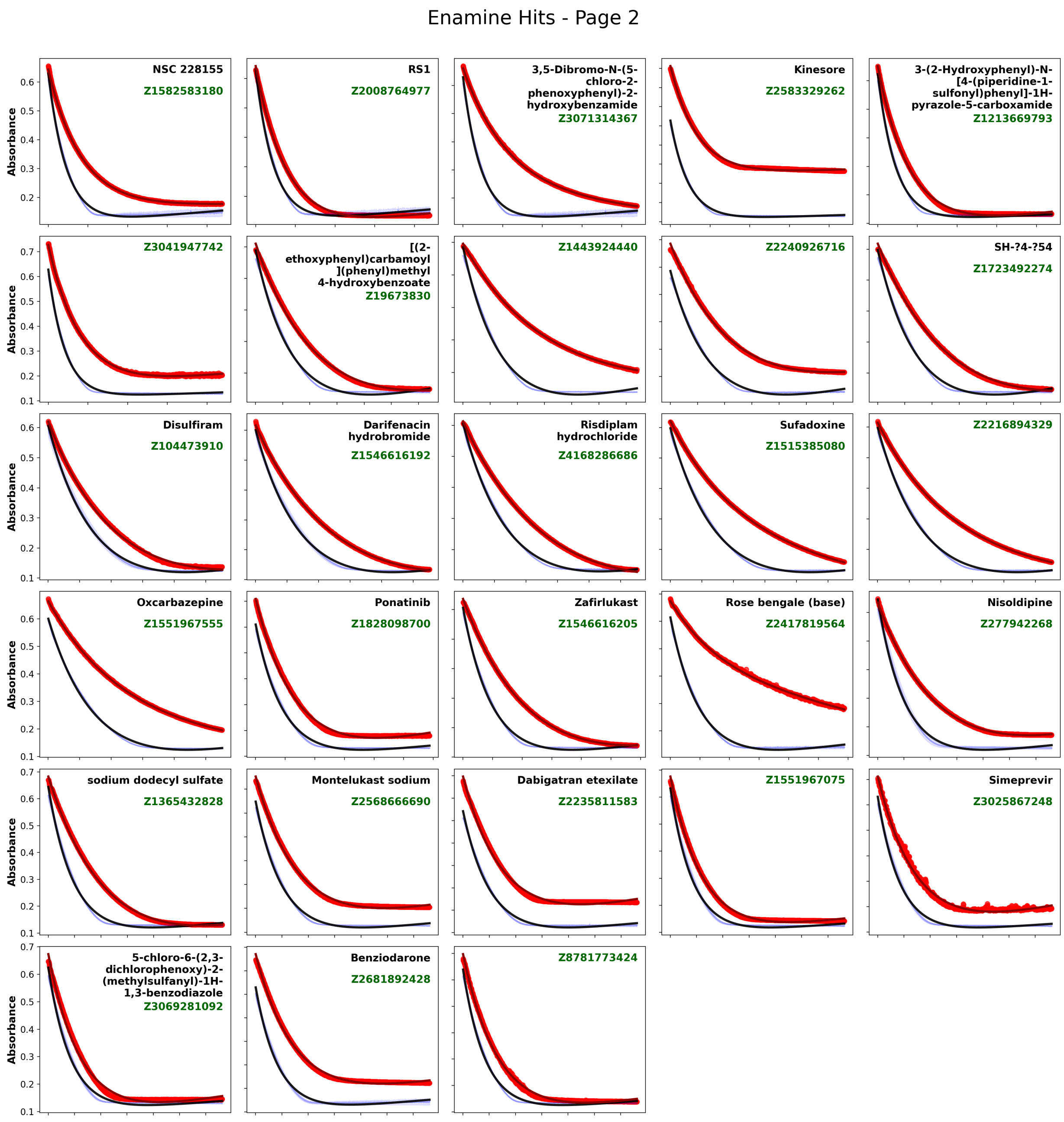


**Figure S2.** Absorbance is measured at 340nm, showing the effect of compounds (red) against DMSO controls (black) in both the LOPAC1280 and the Enamine Bioreference compound libraries. The top right shows the general compound name (if available), and below in green shows the identification number associated with the given library.

**A** NQR-enriched IMV LOPAC1280 hits

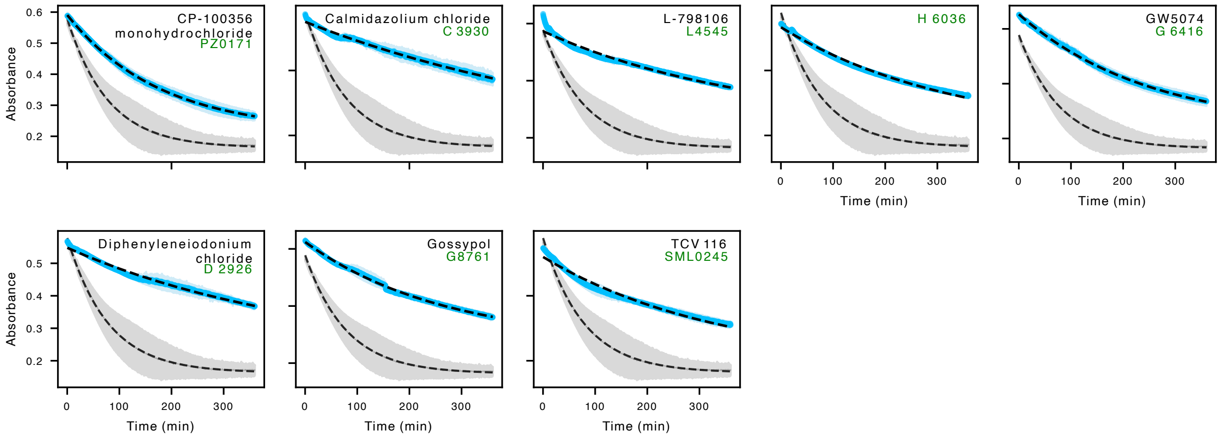


**B** NQR-enriched IMV ENAMINE hits

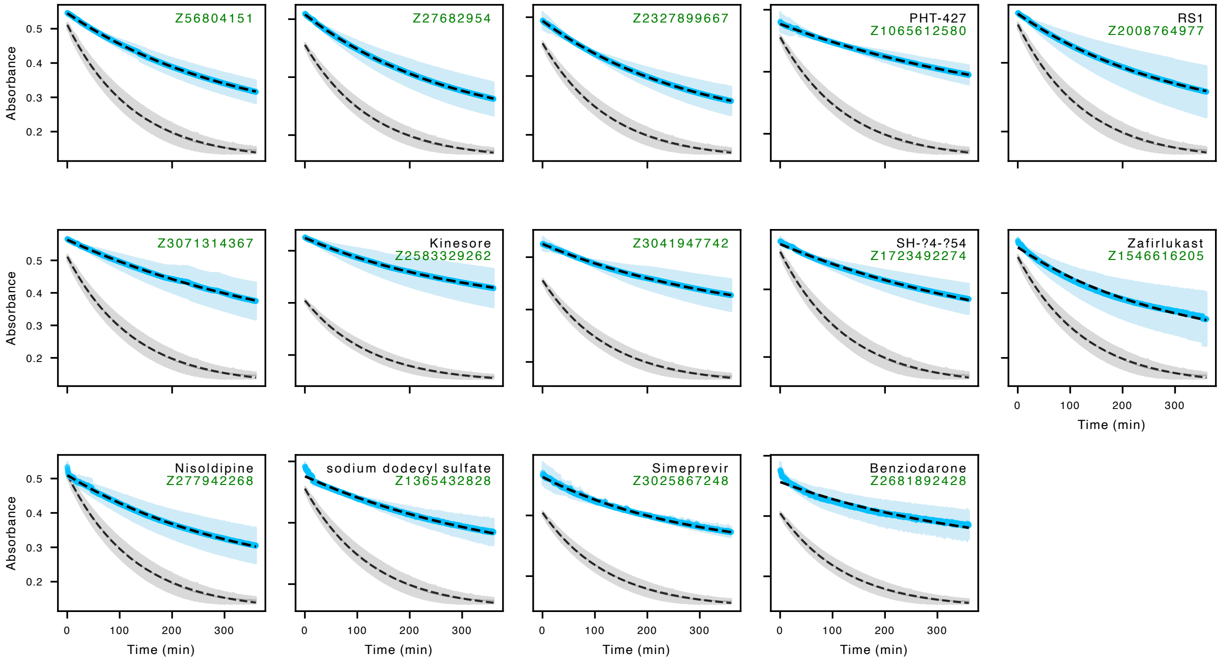


**Figure S3.** Absorbance is measured at 340nm with DMSO control (grey) and the compound (blue) screened at 15µM with *pa*NQR-enriched membranes to verify the target. Top right corner shows the common name of the compound (black, if available) and the identification numbers associated with the drug library (green). (A) shows the verified hits from the LOPAC1280 library, and (B) shows the Enamine Bioreference library.

#### LOPAC1280 *pa*NQR inhibitor structures

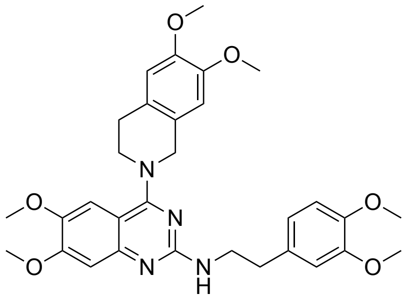

CP-100356
monohydrochloride
PZ0171

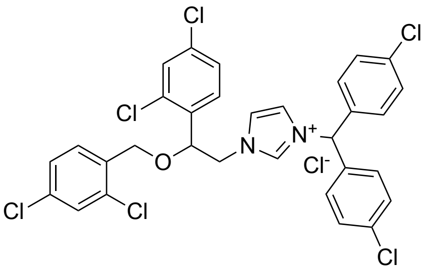

Calmidazolium chloride
C 3930

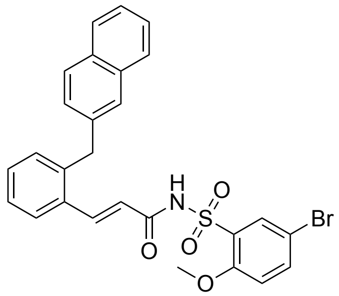

L-798106
L4545

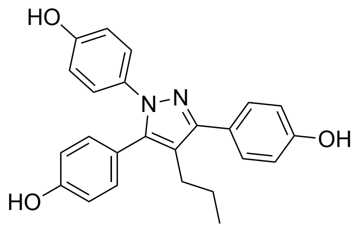

H 6036

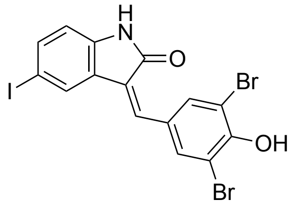

GW5074
G 6416

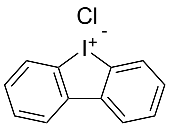

Diphenyleneiodonium
chloride
D 2926

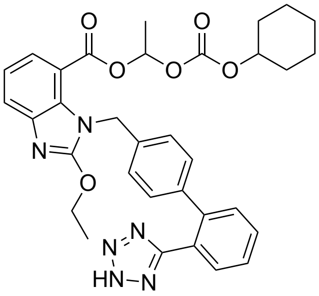

TCV 116
SML0245

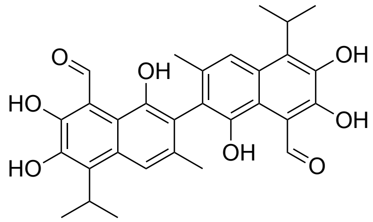

Gossypol
G8761

### ENAMINE bioreference library *pa*NQR inhibitor structures

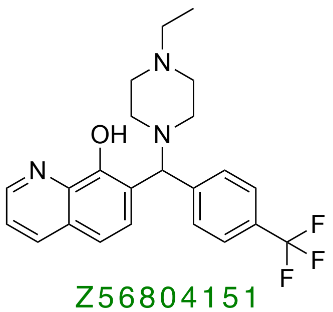

Z56804151

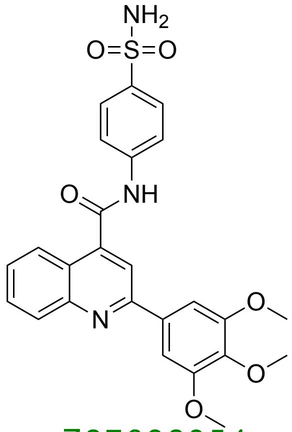

Z27682954

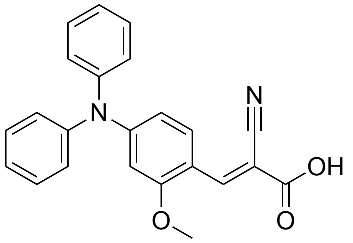

Z2327899667

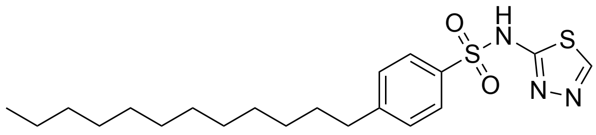

PHT-427
Z1065612580

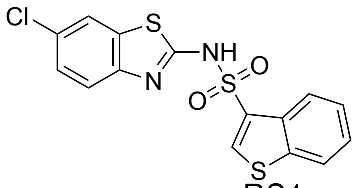

RS1
Z2008764977

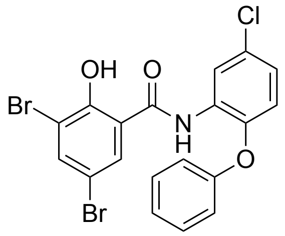

Z3071314367

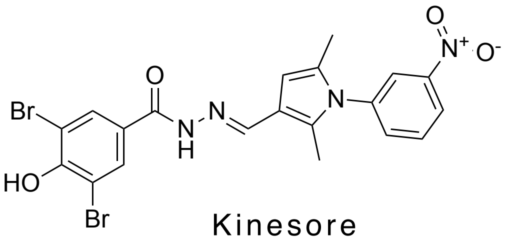

Kinesore
Z2583329262

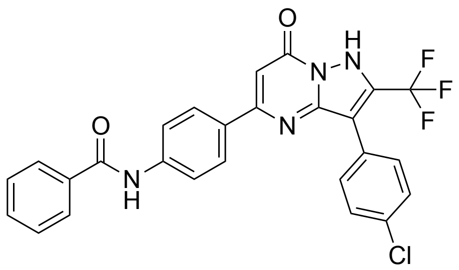

Z3041947742

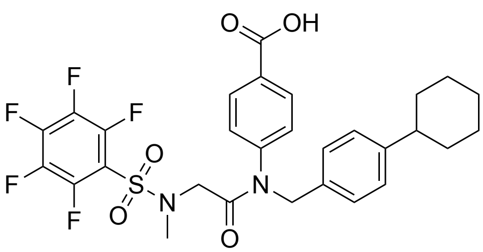

Z1723492274

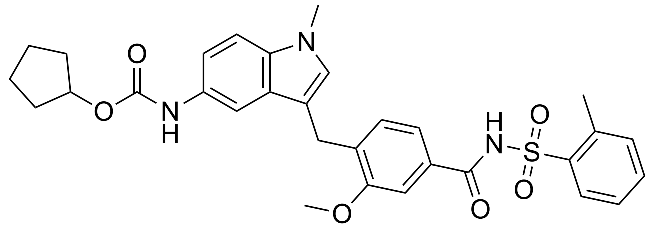

Zafirlukast
Z1546616205

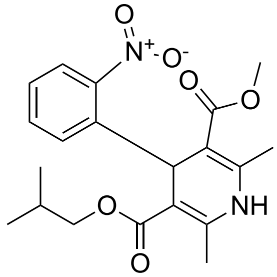

Nisoldipine
Z277942268

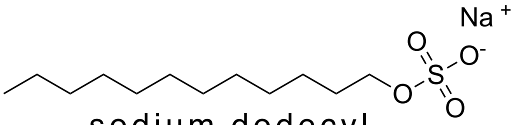

sodium dodecyl sulfate
Z1365432828

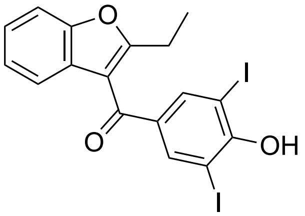

Benziodarone
Z2681892428

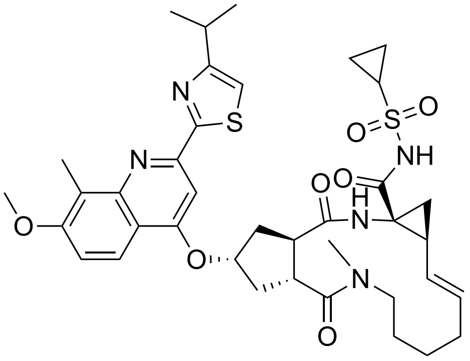

Simeprevir
Z3025867248

**Figure S4.** Structures of hits from the enriched membrane NADH consumption assays. Compounds indicated are from the LOPAC 1280 and the Enamine Bioreference libraries, respectively. The common names for the compounds (black, if available) and the identification numbers associated with the library (green) are given.

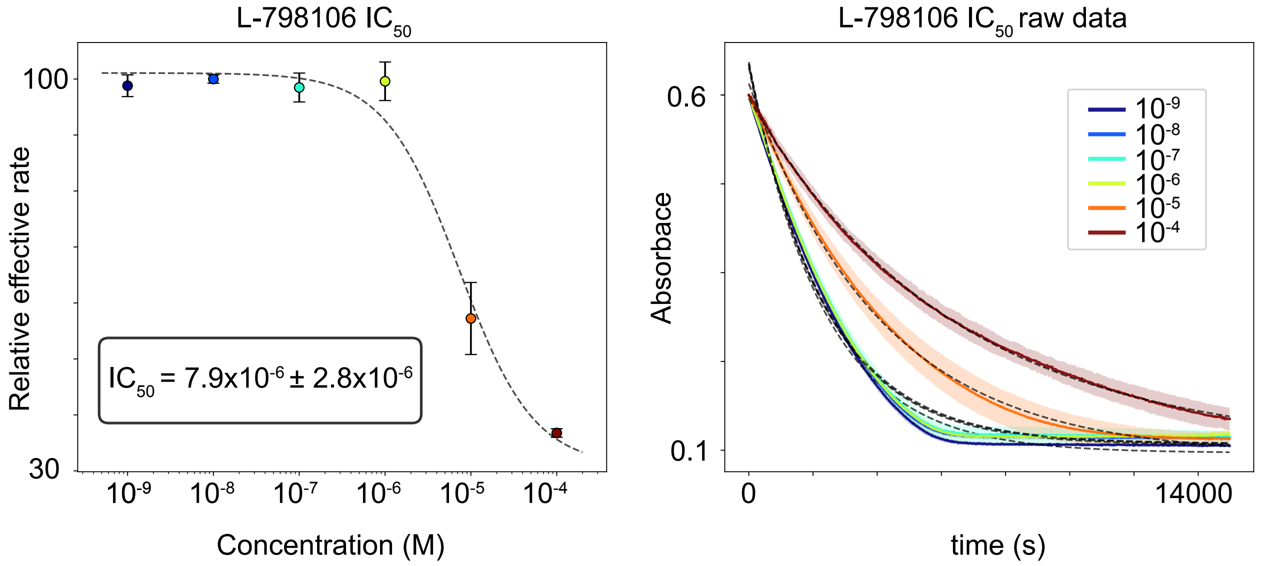


**Figure S5.** On the left is a graph of the dosage dependence curve, and on the right is the raw curve for the NADH oxidation assay with absorbance measured at 340nm. The different colours are associated with different concentrations of the compound in both plots.

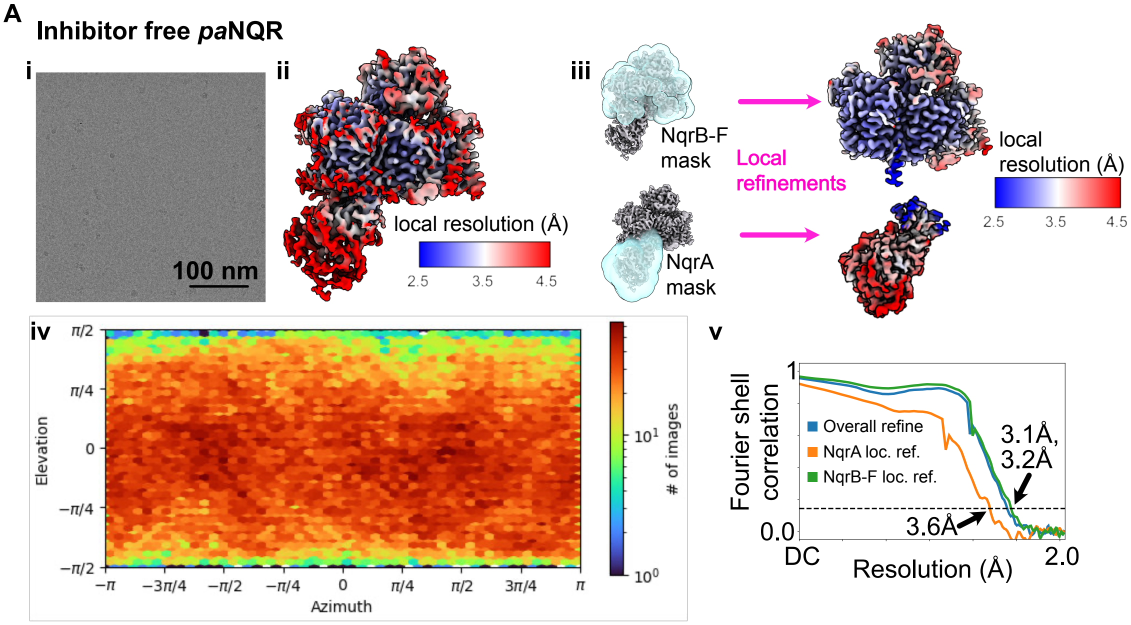


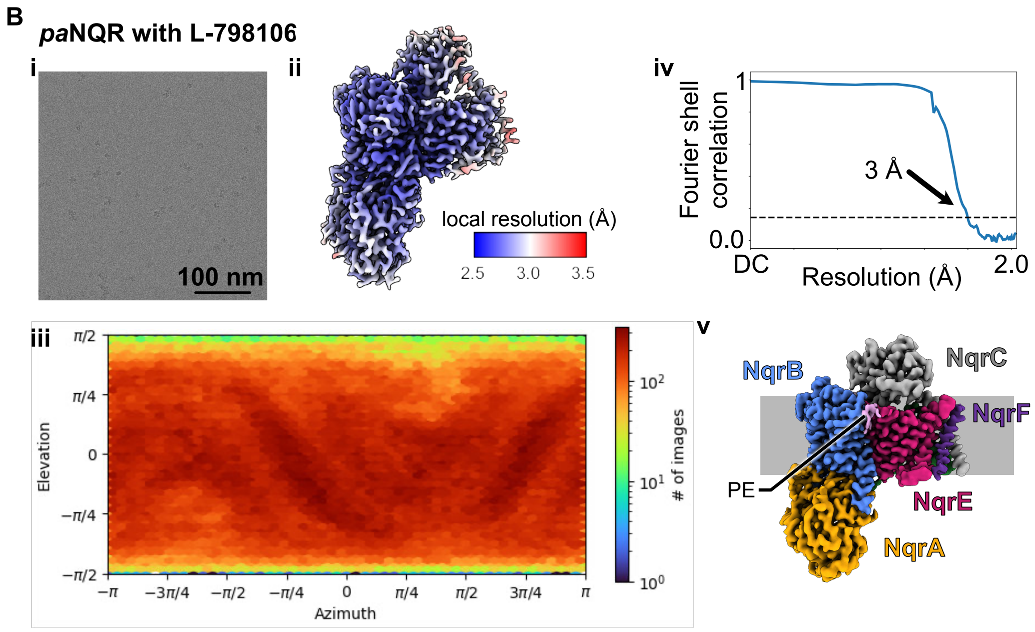


**Figure S6.** Local resolution, viewing direction distribution and fourier shell correlation for inhibitor-free (A) and L-798106-bound (B) datasets.

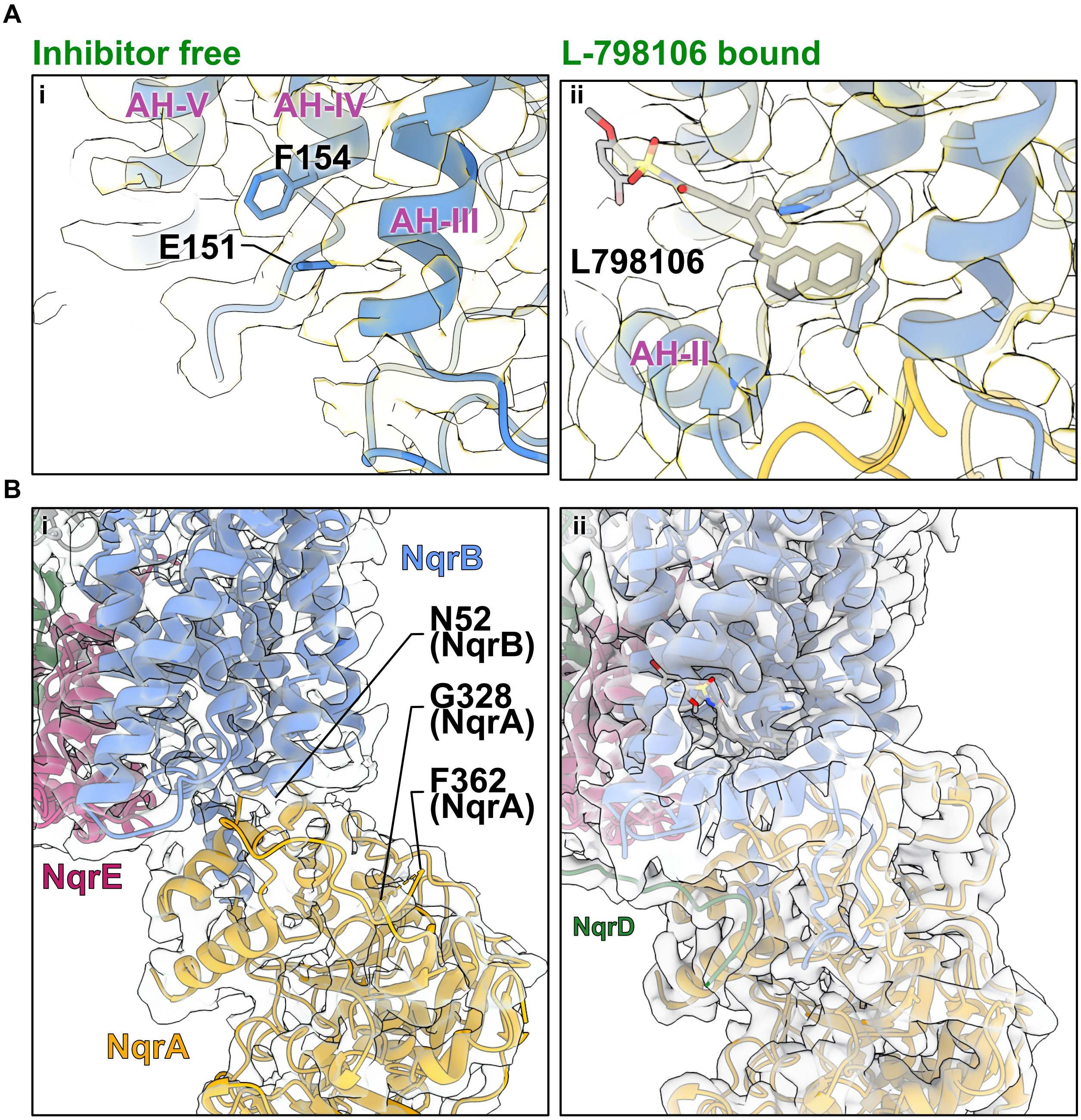


**Figure S7.** Comparison of binding pocket (A) and NqrA and NqrB (B) for inhibitor free (left) and L-798106-bound (right) datasets.

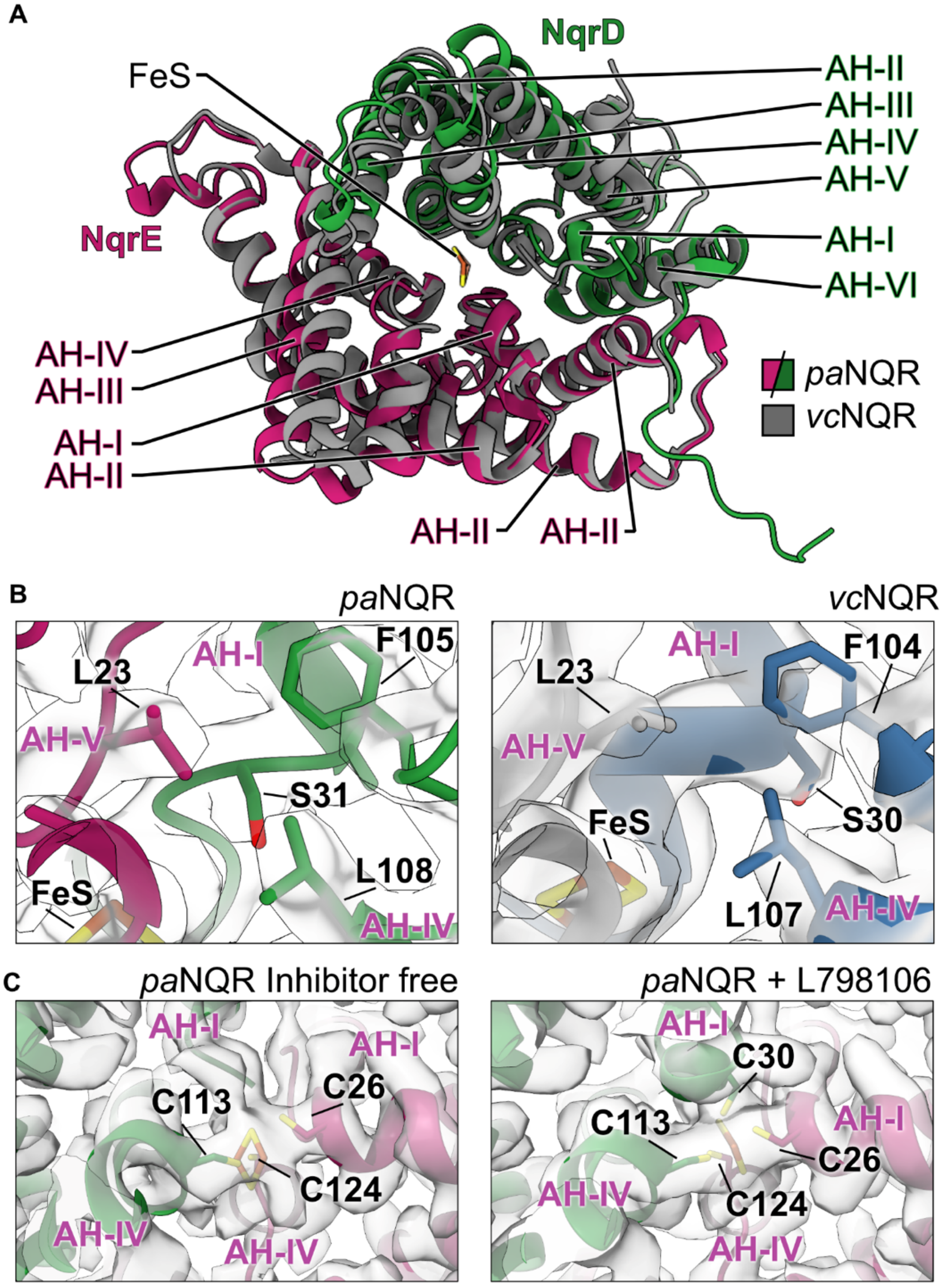


**Figure S8.** (A) Overlay of *vc*NQR (PDB:8EVU) and *pa*NQR NqrE and NqrD subunits. (B) Area around S31 (S30 in *vc*NQR) for both structures with respective maps (EMDB:28637 for *vc*NQR). (C) Maps and models for inhibitor-free and L-798106-bound structures in the area of the iron sulphur cluster.

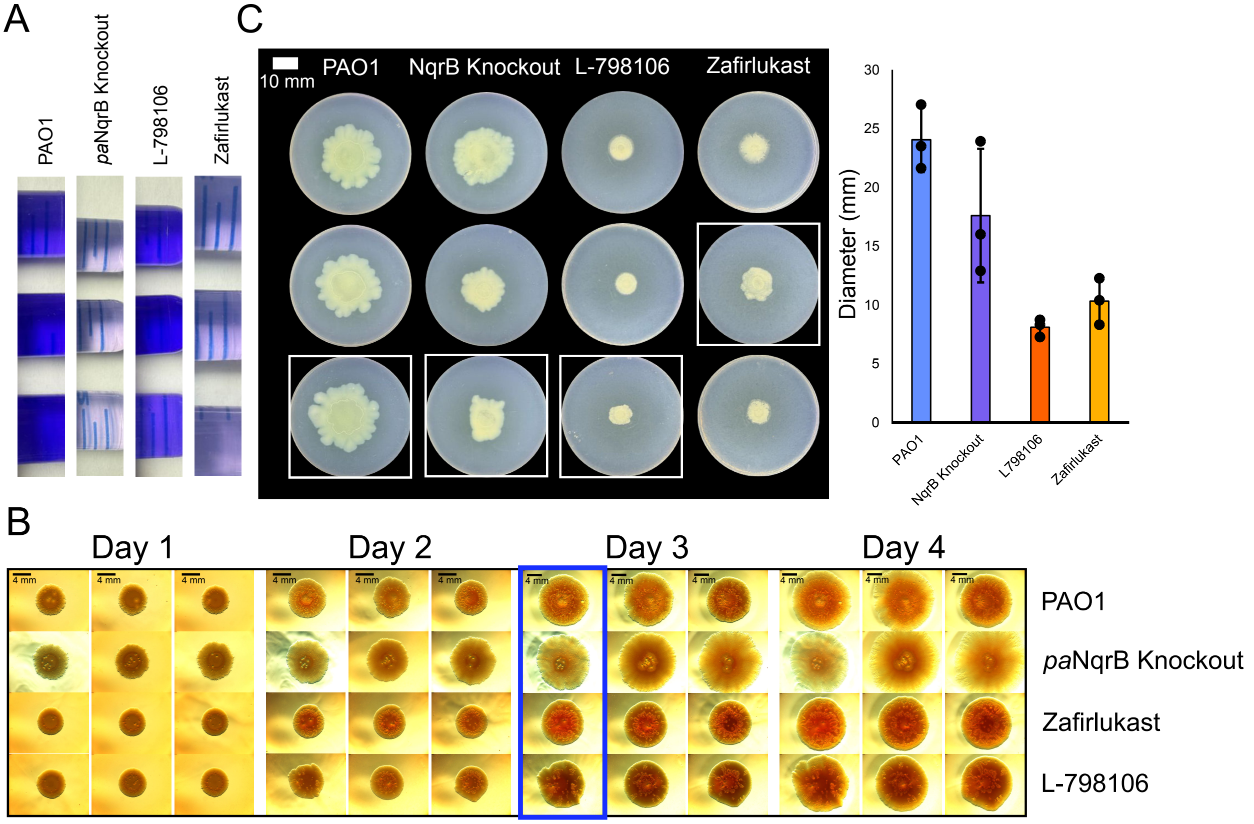


**Figure S9.** (A) Triplicates of crystal violet-stained biofilms resolubilized in a 4:1 mixture of ethanol to acetone, respectively, with compound concentration at 15µM. (B) Congo red biofilm development assay using 1.5% agar in triplicate using PAO1 with and without compound concentrations at 15µM along with ΔNqrB strain, boxed panel indicates replicates chosen for main figure 4C, scale bar is shown in top right. (C) Diameter of the swarming motility colonies in triplicate. (D) Swarming motility assay in triplicate of PAO1 strain with and without compounds at 15µM, along with the ΔNqrB strain in triplicate, boxed plates indicate replicates chosen for main figure 4D, scale bar for each column is shown on the top right.

**Table S1. Bacterial strains and plasmids used.**

| Strain | Description | Reference |
|---|---|---|
| ***E. coli*** | | |
| TOP10 | Cloning strain: F⁻ *mcrA*Δ(*mrr-hsdRMS-mcrBC*) Φ80*lacZ* ΔM15 Δ*lacX74* Δ*araD139*Δ(*ara leu*)7697 *galU galK rpsL endA*1 *nupG*, $Str^R$ | |
| SM10(λpir) | Biparental mating donor strain; *thi thr leu tonA labY supE recA*::RP4-2-Tc::Mu Km λpir, $Kan^R$ | |
| ***P. aeruginosa*** | | **Reference** |
| PAO1 | Wild-type strain | |
| PW2998 | ΔNqrB strain | (Jacobs et al., 2003) |
| **Plasmid** | **Description** | **Reference** |
| **Allelic exchange** | | |
| pEX18Gm | Suicide vector for allelic exchange in *P. aeruginosa*, encodes SacB, $Gen^R$ | (Hmelo et al., 2015) |
| pEX18Gm::PAJDT2 | pEX18Gm with C-terminally 3x Flag tagged *P. aeruginosa* PAO1 *PA2994* cloned between EcorI and HindIII sites, $Gen^R$ | This study |

**Table S2. Cryo-EM data acquisition and image processing.**

| Dataset | Inhibitor free NQR | L-798106-bound NQR |
|---|---|---|
| Electron Microscope | Titan Krios G3 | Titan Krios G4 |
| Camera | Falcon 4i | Falcon 4i |
| Voltage (kV) | 300 | 300 |
| Nominal Magnification | 130,000× | |
| Calibrated physical pixel size (Å) | 0.93 | |
| Total exposure (e/Å$^2$) | 40 | |
| Exposure rate (e/pixel/s) | 4.3 | |
| Number of frames | 29 | |
| Defocus range (µm) | -0.9 to -2 | |
| **Image Processing** | | |
| Motion correction software | *MotionCor2* | *MotionCor2* |
| CTF estimation software | *cryoSPARC v4* | *cryoSPARC v4* |
| Particle selection software | *cryoSPARC v4* | *cryoSPARC v4* |
| Micrographs used | 8,388 | 9,826 |
| Particle images selected | 79,904 | 452,841 |
| 3D map classification and refinement software | *cryoSPARC v4* | *cryoSPARC v4* |

**Table S3. CryoEM map and atomic model statistics.**

| Dataset | Inhibitor free NQR | L-798106-bound NQR |
|---|---|---|
| Modelling and refinement software | Coot, phenix, ISOLDE | Coot, phenix, ISOLDE |
| Protein residues | 1334 | 1502 |
| Ligand | PYT: 1, LMT: 1, FMN: 2, RBF: 1, FES: 1 | PYT: 2, LMT: 1, LIG: 1, FMN: 2, RBF: 1, FES: 1 |
| RMSD bond length (Å) | 0.008 | 0.004 |
| RMSD bond angle (°) | 0.647 | 0.582 |
| Ramachandran outliers (%) | 0 | 0 |
| Ramachandran favoured (%) | 97.48 | 97.31 |
| Rotamer outliers (%) | 0.61 | 0.84 |
| Clash score | 2.55 | 2.77 |
| MolProbability score | 1.14 | 1.20 |
| EMringer score | 3.24 | 3.40 |
